# Draft Genomes of Four *Kluyveromyces marxianus* Isolates Retrieved from the Elaboration Process of Henequén (*Agave fourcroydes*) Mezcal

**DOI:** 10.1101/2023.10.03.560785

**Authors:** Luis Lozano-Aguirre, Morena Avitia, Patricia Lappe-Oliveras, Cuauhtémoc Licona-Cassani, Miguel Ángel Cevallos, Sylvie Le Borgne

## Abstract

We report the draft genomes of four *Kluyveromyces marxianus* isolates obtained from the elaboration process of henequén (*Agave fourcroydes*) mezcal, a Mexican alcoholic beverage. The average nucleotide identity (ANI) analysis revealed that isolates derived from agave plants are distinct from those from other environments, including agave fermentations.

## Announcement

*Kluyveromyces marxianus* is a thermotolerant yeast with a fast growth rate and the ability to metabolize a wide range of carbohydrates, including lactose and inulin in addition to glucose, making it a promising cell factory for industrial biotechnology (1–3). *K. marxianus* has been frequently isolated from dairy products (4) and a variety of other habitats such as fermented beverages (5), plants and fruits (6, 7), sugarcane mills (8, 9) as well as clinical samples among others. It has been suggested that isolates from agave and associated fermentations may constitute a new clade within the *K. marxianus* species (4). Among the twenty-one *K. marxianus* genomes available in the NCBI genome database only two correspond to strains derived from agave; strains UFS-Y2791 from an agave plant (unpublished) and SLP1 from spontaneous mezcal fermentation (5). Here, we present the draft genome sequences of four *K. marxianus* isolates retrieved from the elaboration process of henequén mezcal, a Mexican fermented alcoholic beverage elaborated from the juice extracted from cooked henequén cores (10). Henequén (*Agave fourcroydes*) is an agave species native to the Yucatan peninsula.

DNA was prepared from overnight cultures in yeast extract-peptone-dextrose (YPD) broth at 30°C using the Quick-DNA Fungal/Bacterial Miniprep Kit (Zymo Research) following the manufacturer’s instructions. DNA quality and purity were assessed by 0.7% (w/v) agarose gel electrophoresis in 1x TBE buffer and UV absorbance measurements on a Nanodrop 2000 spectrophotometer (Thermo Scientific). DNA was quantified using a Qubit 3.0 fluorometer (Life Technologies). Paired-end genomic DNA libraries were constructed using the TruSeq Nano kit (Illumina) according to the manufacturers’ instructions. Libraries quality and quantity were verified using a 2100 BioAnalyzer (Agilent Technologies). Sequencing was performed on the Illumina HiSeq 2500 platform through the standard rapid-sequencing protocol to generate 150-bp paired-end reads.

Reads quality was assessed with FastQC v0.11.9 (11). The adapters and low-quality bases were discarded using Trimmomatic v.0.39 by performing sliding window trimming with a minimum average quality of 20 and a minimum sequence read length of 36 bases (12). *De novo* genome assemblies were generated using Velvet v.1.2.10 (13) and Spades v.3.12.0 (14) and the obtained assemblies were merged using Metassembler v.1.5 (15). Assemblies quality was assessed using QUAST v.4.1 (16). Gene prediction and functional annotation was performed with Funannotate (v.1.8.14) using *Kluyveromyces lactis* as the training species (17). Assemblies completeness was evaluated with BUSCO v.5.4.7 using the saccharomycetes_odb10 database (18). Average nucleotide identity (ANI) analysis was calculated with pyani v0.2 (19). The heatmaps were built with the R packages ggplot2 and pheatmap. Table 1 details the sequencing data, assemblies statistics, BUSCO scores and ANI values.

**TABLE 1.** General features of the sequenced genomes.

| Isolate | Origin | No. of reads | No. of contigs | Total length (Mb) | Coverage (x) | GC content (%) | N50 (kb) | L50 (kb) | BUSCO score (%) | ANI score (%)<sup>a</sup> | ANI coverage (%)<sup>a</sup> | No. of predicted genes | No. of proteins | Assembly accession no. | SRA accession no. |
| --- | --- | --- | --- | --- | --- | --- | --- | --- | --- | --- | --- | --- | --- | --- | --- |
| Kmx14 | Henequén leaf | 4,395,815 | 215 | 10.6 | 116 | 39.99 | 87,645 | 37 | 98.9 | 94.5 | 94.9 | 4958 | 4781 | SAMN31841251 | SRR23105050 |
| Kmx16 | Nonfermented henequén cooked juice | 4,686,443 | 219 | 10.5 | 124 | 40.04 | 87,096 | 36 | 98.7 | 94.6 | 94.2 | 4868 | 4697 | SAMN31841253 | SRR23105048 |
| Kmx22 | Fermented henequén cooked juice | 5,405,732 | 396 | 10.6 | 142 | 40.11 | 45,399 | 74 | 99.4 | 99.2 | 96.9 | 4952 | 4789 | SAMN31841254 | SRR23105047 |
| Kmx24 | Cooked henequén core | 3,024,326 | 218 | 10.6 | 79 | 39.99 | 78,647 | 40 | 99.1 | 94.5 | 95 | 4899 | 4721 | SAMN31841255 | SRR23105046 |

<sup>a</sup> ANI against the reference genome *K. marxianus* DMKU3-1042

According to the ANI analysis (Figure 1), isolates UFS-Y2791, Kmx14, Kmx16 and Kmx24, from agave plant, henequén leaf, nonfermented henequén cooked juice and cooked henequén core formed a separate group with ANI values greater than 99% between each other. Interestingly, strains Kmx22 and SLP1 from fermented henequén cooked juice and mezcal fermentations, respectively, did not belong to this group and exhibited more relatedness to *K. marxianus* isolates from dairy and other environments. These data confirm that there is further yeast diversity to be accessed in agave environments (4) in a similar way to what has been described for cactus-yeasts (20) .

**Figure 1.**
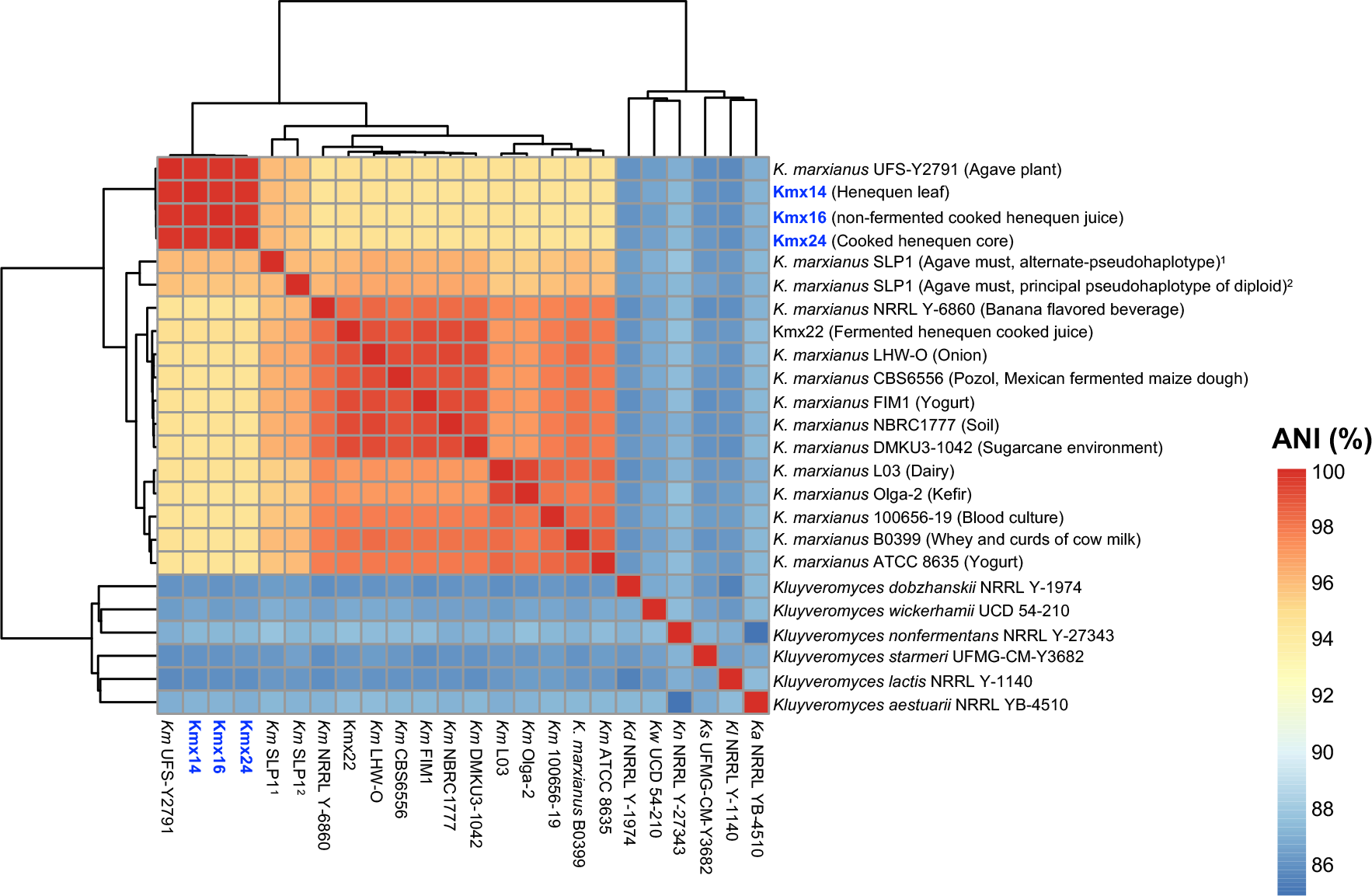
Heatmap of ANI values. The isolates sequenced here are indicated in blue color. The *Kluyveromyces* genomes were downloaded from NCBI: *K. marxianus* UFS-Y2791 (GenBank accession number GCA_001692465.1), *K. marxianus* SLP1 alternate-pseudohaplotype (GCA_021014395.1), *K. marxianus* SLP1 principal pseudohaplotype of diploid (GCA_021014425.1), *K. marxianus* NRRL Y-6860 (GCA_002356615.1), *K. marxianus* LHW-O (GCA_003046155.1), *K. marxianus* CBS6556 (GCA_016625955.1), *K. marxianus* FIM1 (GCA_001854445.2), *K. marxianus* NBRC 1777 (GCA_001417835.1), *K. marxianus* DMKU3-1042 (GCA_001417885.1), *K. marxianus* L03 (GCA_008000265.1), *K. marxianus* Olga-2 (GCA_016584165.1), *K. marxianus* 100656-19 (GCA_902364165.1), *K. marxianus* B0399 (GCA_001660455.1), *K. marxianus* ATCC 8635 (GCA_017309885.1), *K. dobzhanskii* NRRL Y-1974 (GCA_003705805.2), *K. wickerhamii* UCD 54-210 (GCA_000179415.1), *K. nonfermentans* NRRL Y-27343 (GCA_003670155.1), *K. starmeri* UFMG-CM-Y3682 (GCA_008973615.1), *K. lactis* NRRL Y-1140 (GCA_000002515.1) and *K. aestuarii* NRRL YB-4510 (GCA_003707555.1).

## Data availability statement

Assemblies are available at the National Center for Biotechnology Information (NCBI) with assembly accession numbers indicated in Table 1. The genome assembly generated in this study and the reads are deposited under BioProject ID PRJNA904382.

## Acknowledgments

This research was supported by the Universidad Autónoma Metropolitana-Unidad Cuajimalpa in Mexico City (Research project 87 S210-21 “Caracterización y potencial de aplicación de levaduras y bacterias autóctonas de México, DCNI-05-210-21) and the CONAHCyT research grant CB-2010-01 156451 awarded to Sylvie Le Borgne.

## Notes

### Competing Interest Statement

The authors have declared no competing interest.

